# Experimental and computational studies on molecular mechanism by which Curcumin allosterically inhibits Dengue protease

**DOI:** 10.1101/2020.09.20.305664

**Authors:** Liangzhong Lim, Mei Dang, Amrita Roy, Jian Kang, Jianxing Song

## Abstract

Flaviviruses including DENV and ZIKV encode a unique two-component NS2B-NS3 protease essential for maturation/infectivity, thus representing a key target for designing anti-flavivirus drugs. Here for the first time, by NMR and molecular docking, we reveal that Curcumin allosterically inhibits the Dengue protease by binding to a cavity with no overlap with the active site. Further molecular dynamics (MD) simulations decode that the binding of Curcumin leads to unfolding/displacing the characteristic β-hairpin of the C-terminal NS2B and consequently disrupting the closed (active) conformation of the protease. Our study identified a cavity most likely conserved in all flaviviral NS2B-NS3 proteases, which could thus serve as a therapeutic target for discovery/design of small molecule allosteric inhibitors. Moreover, as Curcumin has been used as a food additive for thousands of years in many counties, it can be directly utilized to fight the flaviviral infections and as a promising starting for further design of potent allosteric inhibitors.

## INTRODUCTION

Dengue virus (DENV) of the *Flaviviridae* family, which also includes several other human pathogens such as Zika (ZIKV), West Nile (WNV), Japanese encephalitis and Yellow fever viruses, is the most prevalent human pathogens transmitted by *Aedes* mosquitoes with 3.6 billion people at risk, particularly in tropical and subtropical regions. Annually over 390 million human infections occur in ∼110 countries including the southern US and Singapore which leads to ∼25,000 deaths mostly among children (1-4). DENV causes dengue fever, dengue haemorrhagic fever and dengue shock syndrome. Despite intense studies, so far, no marketed antiviral drug exists to effectively treat dengue associated diseases (4-6).

The DENV genome is composed of an 11-kb single-stranded positive sense RNA, which is translated into a large polyprotein by the host-cell machinery upon infection. The polyprotein of the *Flaviviridae* family needs to be subsequently processed into 10 proteins, which include three structural proteins (capsid, membrane, and envelope) and seven nonstructural proteins (NS1, NS2A/B, NS3, NS4A/B, and NS5). The structural proteins constitute the viral particle while the nonstructural proteins are involved in the replication of the RNA genome, virion assembly, and attenuation of the host antiviral response, thus essential for replication of all flaviviruses. The correct processing of the polyprotein is implemented by host cell proteases including furin and signalaseas, as well as a virus-encoded NS2B-NS3 protease, which thus has been established as a valuable target for drug design to treat DENV and other flavivirus infections (4-24).

The Dengue protease domain consists of the N-terminal part of the NS3 protein adopts a chymotrypsin-like fold consisting of two β-barrels, each composed of six β-strands, with the catalytic triad (His51-Asp75-Ser135) located at the cleft between the two β-barrels (Fig. 1A). Unlike other proteases with a chymotrypsin-like fold, the flavivirus proteases including dengue one, additionally require a stretch of ∼40 amino acids from the cytosolic domain of NS2B for its catalytic function, thus called a two-component protease. While the protease domains adopt highly similar structures in all crystal structures, the NS2B cofactor was found to assume two distinctive structures, namely the inactive or open form in the unbound state (I of Fig. 1A) as well as active or closed form in complex with the substrate peptide (II of Fig. 1A) by X-ray crystallography (8-11). Furthermore, recent NMR studies revealed that in solution, the Dengue NS2B-NS3 protease is very dynamic and undergoes the exchange between two conformations. However, the closed conformation is the major form even in the unbound state, which thus represents the best model for structure-guided drug designs (12-15).

**Figure 1.**
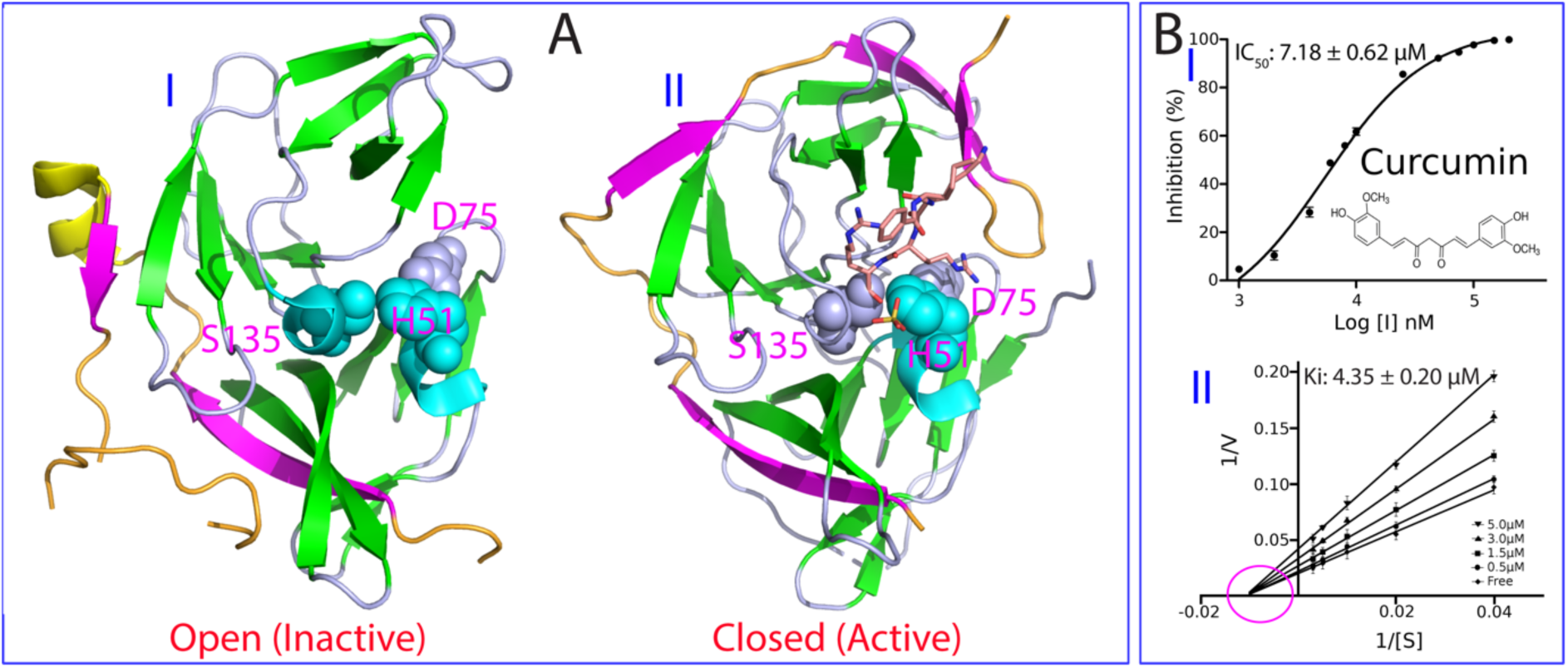
Curcumin inhibits the Dengue NS2B-NS3 protease in a non-competitive mode. (A) Crystal structures of the Dengue NS2B-NS3 protease in the open (inactive) conformation (PDB ID of 2FOM) in the unbound state (I); and in the closed (active) conformation (PDB ID of 3U1I) in complex with a substrate peptide in sticks (II). The β-strand is colored in purple, helix in yellow and loop in brown for the NS2B cofactor, while the β-strand is in green, helix in cyan and loop in light blue for the NS3 protease domain. The catalytic triad His51-Asp75-Ser135 are displayed in spheres and labeled. (B) Chemical structure of Curcumin and the inhibitory data used for fitting IC50 value for Curcumin (I). Lineweaver-Burk plot for determining inhibitory constant (Ki) of Curcumin on the Dengue NS2B-NS3 protease (II). [S] is the substrate concentration; v is the initial reaction rate. The curves were generated by the program GraphPad Prism 7.0. The purple circle is used to indicate that the inhibition is non-competitive, characteristic of the same Km but varying Vmax values in the presence of Curcumin at different concentrations.

Previous efforts for drug development targeting the flaviviral NS2B-NS3 proteases revealed the major challenge in rational design of their active site inhibitors: their active sites are relatively flat (9-24). In this context, to respond to the urgency to fight ZIKV and DENV infection in Singapore, previously we conducted an intense attempt to screen inhibitors for the Zika and Dengue NS2B-NS3 proteases from natural products isolated from edible plants. We successfully identified a natural phenol Curcumin, or 1,7-bis(4-hydroxy-3 methoxyphenyl)-1,6-heptadiene-3,5-dione (Fig. 1B), with a significant inhibitory effect on both proteases (21). Further analysis of enzymatic kinetics revealed that Curcumin inhibits the Zika NS2B-NS3 protease in a non-competitive mode. In other words, Curcumin may act as an allosteric inhibitor for the NS2B-NS3 proteases.

Curcumin is of both fundamental and therapeutic interest because it has a significant inhibitory effect on the Zika NS2B-NS3 protease (IC50 of 3.45 μM and Ki of 2.61 μM) as we previously determined (21). Very recently, Curcumin and its four analogues were also found to inhibit the Dengue NS2B-NS3 protease as well as replicon replication in DENV-infected cells (4). Moreover, Curcumin is isolated from a very popular food additive yellow ginger Turmeric (*Curcuma longa*). A huge number of previous studies have shown that Curcumin owns a diversified biological and pharmaceutical activities. including antitumoral, antimicrobial, anti-inflammatory, antioxidant, antihepatotoxic, antihyperlipidemic, antiviral, and anti-Alzheimer’s disease effects. (4,25-27) In this study, we aimed to understand the mechanism by which Curcumin inhibits the Dengue NS2B-NS3 protease with biochemical assay and biophysical methods including NMR spectroscopy and molecular dynamics (MD) simulations. For the first time, our NMR studies reveal that in contrast to the active-site inhibitors which act to reduce the dynamics of the Dengue NS2B-NS3 protease (15), Curcumin significantly increased the backbone dynamics of the Dengue protease particularly on µs-ms time scale. Nevertheless, we have successfully established the binding mode as derived from the NMR-derived constraints, showing that Curcumin binds to a cavity of the Dengue NS2B-NS3 protease which has no overlap with its active site. Further MD simulations reveal that the binding of Curcumin leads to the disruption of the closed conformation which is essential for its catalytic activities. Altogether, our study provides a dynamic view of the mechanism by which Curcumin allosterically inhibit the Dengue NS2B-NS3 protease through mediating the equilibrium between the open and closed conformations. Therefore, the modulation of this conformational equilibrium might indeed represent a promising strategy to discover/design small molecules for allosterically inhibiting the flaviviral NS2B-NS3 proteases to treat flavivirus infections.

## RESULTS

### Inhibition of the Dengue protease by Curcumin

Previous studies have extensively shown that the Dengue NS2B-NS3 protease with the NS2B and NS3 covalently unlinked better represents the *in vivo* state, and also manifested well-dispersed NMR spectra which thus allowed the NMR assignment (14,16). As such, in the present study we used the same unlinked version of the Dengue NS2B-NS3 protease we previously constructed (16) for all experiments. We determined its Km to be 89.39 ± 6.62 μM; and Kcat to be 0.12 ± 0.01 s^-1^, which are almost identical to our previous results with Km = 92.39 ± 9.94 μM and Kcat = 0.15 ± 0.01 s^-1^ (16).

In our previous screening, we found that Curcumin also showed significant inhibitory effect on the Dengue NS2B-NS3 protease. Here, we further determined its value of IC50 to be 7.18 ± 0.62 μM and inhibitory constant Ki to be 4.35 ± 0. 02 μM (Fig. 1B), which are only slightly different from those on the Zika NS2B-NS3 protease (IC50 of 3.45 μM and Ki of 2.61 μM). As we previously observed on the Zika NS2B-NS3 protease (21), Curcumin also inhibited the Dengue NS2B-NS3 protease by changing Vmax but not Km (II of Fig. 1B), thus indicating that Curcumin also acts as a non-competitive inhibitor for the Dengue NS2B-NS3 protease.

### NMR characterization of the binding of Curcumin to the Dengue protease

To gain insights into the binding mode of Curcumin to the Dengue NS2B-NS3 protease, we have successfully obtained the protease samples with either the NS2B cofactor or NS3 protease domain selectively ^15^N-labeled. In the NS2B-NS3 protease complex, both ^15^N-labeled NS2B (Fig. S1A) and NS3 protease domain (Fig. S1B) have well-dispersed HSQC spectra, typical of the well-folded protein. Furthermore, the chemical shifts of their HSQC peaks are very similar to what were previously assigned (14,16).

Very unexpectedly, however, when we titrated the Dengue protease samples with Curcumin at molar ratios of 1:0.5, 1:1; 1:2.5, 1:5 and 1:10 (Protease:Curcumin), no significant shifts were observed for most HSQC peaks of both NS2B (Fig. S1A) and NS3 protease domain (Fig. S1B), indicating that the binding did not induce significant structural changes. On the other hand, the HSQC peaks became step-wise broad and consequently their intensity gradually reduced. At 1:10, many well-dispersed HSQC peaks became too weak to be detectable.

Usually, the line-broadening of HSQC peaks upon binding results from the micro-molar dissociation constants, and/or binding-induced increase of conformational exchanges particularly on µs-ms time scale (12-16,28-31). For example, previously we found that the binding of small molecules to EphA4 receptor with micro-molar Kd led to significant broadening of EphA4 HSQC peaks (28,29). Furthermore, for a well-folded protein, a slight disruption/destabilization of the native structure was sufficient to trigger the significant increase of conformational exchange on µs-ms time scale with many well-dispersed HSQC peaks broadened/disappeared (30,31). Here, the Curcumin binding-provoked increase of conformational exchanges might explain the fact that our ITC measurements of the binding of Curcumin to the Dengue NS2B-NS3 all gave rise to the data with a very high level of noises.

The current results indicate that the binding of Curcumin would lead to significant increase of dynamics particularly on µs-ms time scale, which is in a sharp contrast to a recent NMR study of the interaction of the Dengue NS2B-NS3 protease with an active-site inhibitor, in which the inhibitor binding led to a significantly reduced dynamics, thus manifesting high-quality NMR spectra (15). Consequently, here due to the significant weakening of the intensity of HSQC peaks of the Dengue NS2B-NS3 protease upon binding to Curcumin, despite intense attempts we were unable to acquire high-quality NMR relaxation data to derive the backbone dynamics of the protease bound to Curcumin as we previously performed on other proteins on ps-ns and μs-ms time scales (29,30).

### NMR-guided molecular docking of the Curcumin-binding mode

Due to the significant increase of the protease dynamics upon binding to Curcumin which led to severe NMR line broadening, we were also unable to further determine the structure of the protease complex with Curcumin by NMR spectroscopy. Furthermore, we also intensely attempted to crystalize the complex sample by screening a large array of buffer conditions but all failed. Therefore, here we analysed the intensity of HSQC peaks in the presence of Curcumin at different molar ratios, and Fig. 2 presents the normalized intensity of the NS2B (Fig. 2A) and NS3 protease domain (Fig. 2B) in the presence of Curcumin at 1:5. The NS2B has an average intensity of 0.63 while the NS3 protease domain has an average intensity of 0.76, implying that slightly more dynamics were provoked on the NS2B cofactor than on the NS3 protease domain. Noticeably, some residues have significantly reduced HSQC peak intensity (< average – one STD), which may be involved in the binding to Curcumin. While the 5 significantly-perturbed residues including Arg55, Ala56, Leu74, Ile78 and Gly82 are located over the whole NS2B, the 14 significantly-perturbed residues of the NS3 protease domain including Gly87, W89, Leu115, Phe116, Ly117, Thr118, Asn119, Thr120, Thr122, Ile123, Val162, Ser163, Ala164 and Ile165 are clustered together to form a cavity with on overlap with the catalytic triad (Fig. 2C).

**Figure 2.**
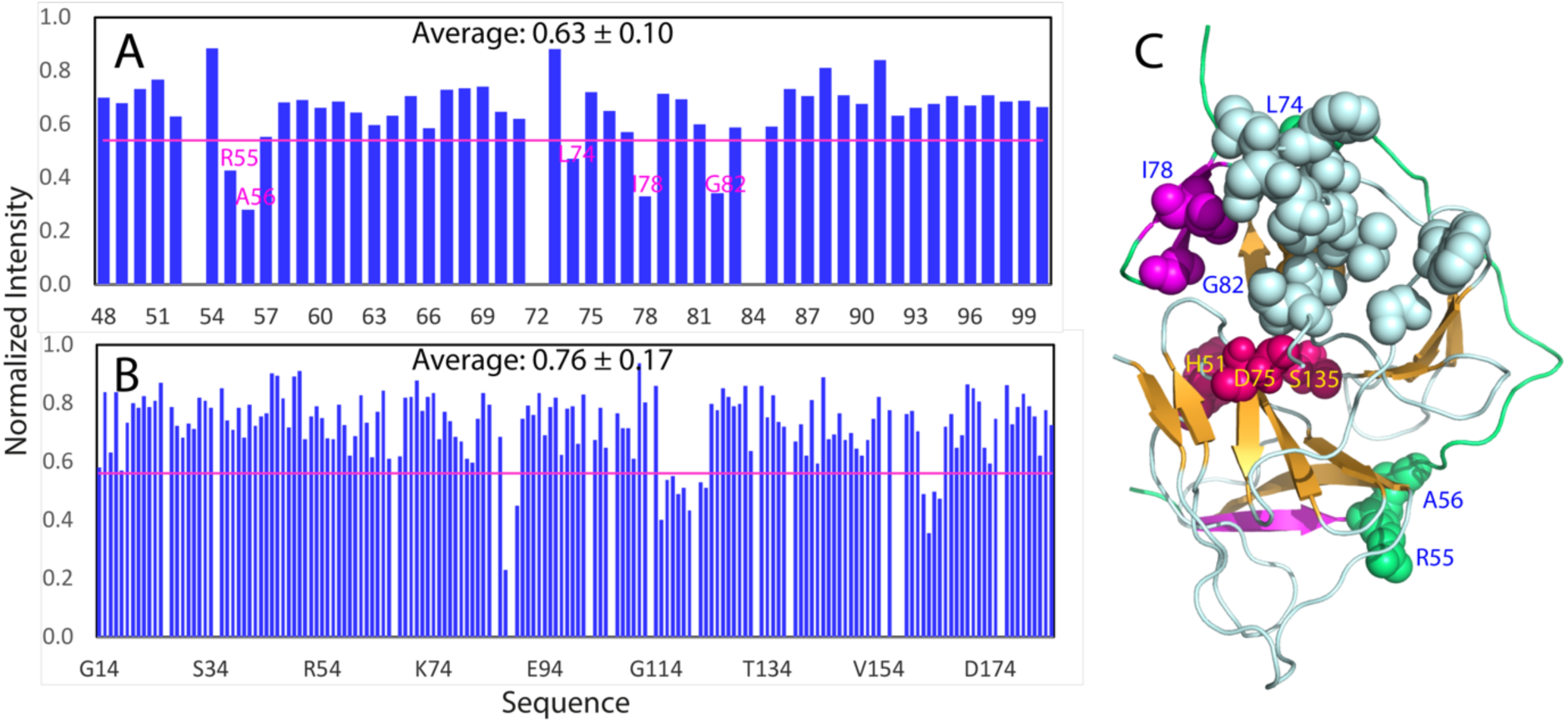
NMR identification of the perturbed residues upon binding Curcumin. (A) Normalized HSQC peak intensity of the selectively ^15^N-labeled NS2B in the complex with unlabeled NS3 in the presence of Curcumin at a molar ratio of 1:5. (B) Normalized HSQC peak intensity of the selectively ^15^N-labeled NS3 in the complex with unlabeled NS2B in the presence of Curcumin at a molar ratio of 1:5. Significantly perturbed residues are defined as those with the normalized intensity < 0.53 for NS2B and < 0.59 for NS3 protease domain (average value - one standard deviation). (C) The structure of the Dengue NS2B-NS3 protease with the significantly perturbed residues displayed in spheres. The significantly perturbed 5 residues within the NS2B cofactor are labeled. The catalytic triad residues His51-Asp75-Ser135 were also displayed in red spheres. The β-strand is colored in purple and loop in green for the NS2B cofactor, while the β-strand is in brown, helix in cyan and loop in light blue for the NS3 protease domain.

With the above-identified 19 residues set as active residues, we constructed the Curcumin-protease complex by the well-established program HADDOCK (32), as we previously utilized to build up the small molecule-protein and ATP-protein complexes (33,34). Fig. 3A presents the complex model with the lowest energy score, in which Curcumin inserts into a cavity constituted by the residues from both NS2B and NS3 protease domain, mainly including those identified by NMR titrations (I of Fig. 3A). Indeed, this cavity has no direct overlap with the catalytic triad, suggesting that Curcumin allosterically inhibits the Dengue NS2B-NS3 protease, which is completely consistent with the inhibitory profile of Curcumin (II of Fig. 1B). It is worthwhile to note that this cavity is close to Ala125, whose covalent modification has been previously identified to allosterically inactivate the catalytic activity of the protease (18). Therefore, we also docked Curcumin in both keto-and enol-forms to the Dengue protease with Ala125 modified in the open form (PDB ID of 4M9T) using the same active residues. As shown in III of Fig. 3A, Curcumin in both forms also bind to the very similar cavity of the Dengue protease in the open form, indicating that this site also exists in the open (inactive) form.

**Figure 3.**
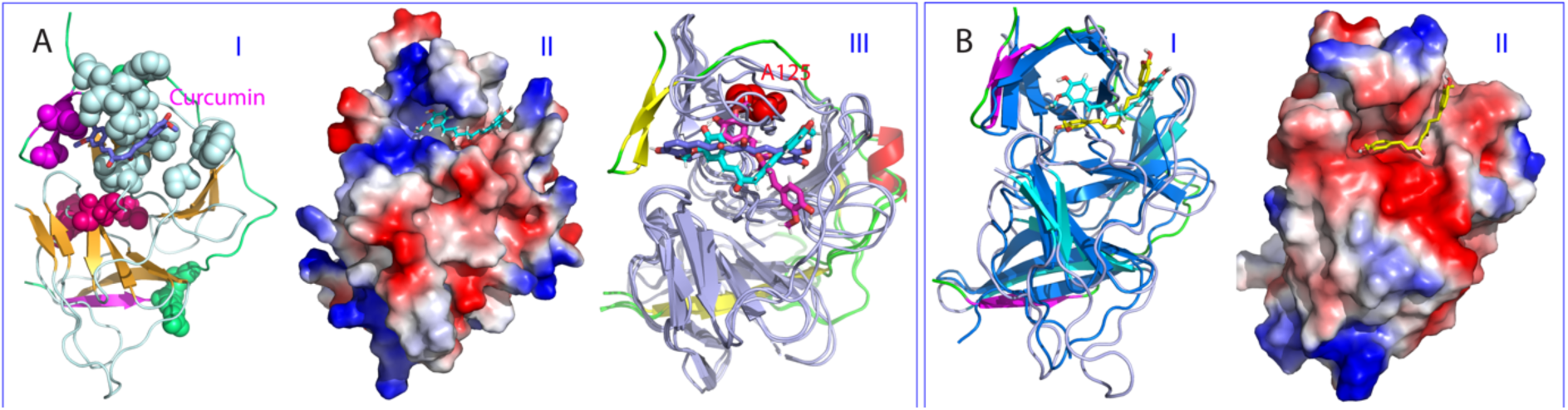
Docking structures of the Curcumin-protease complexes. (A) Docking structure of the Dengue NS2B-NS3 protease in complex with Curcumin in ribbon with the β-strand colored in purple and loop in green for the NS2B cofactor, as well as the β-strand in brown, helix in cyan and loop in light blue for the NS3 protease domain (I); and in electrostatic potential surface (II). (III) Superimposition of the complexes of Curcumin with the Dengue NS2B-NS3 protease in the closed and open states. The NS2B is colored as: yellow for *β*-strand, red for *α*-helix and green for loop, while NS3 is colored in bright blue. Cyan is for Curcumin complexed with the protease in the closed state while blue for Curcumin (enol-form) and pink for Curcumin (keto-form) complexed with the protease in the open state (PDB ID of 4M9T). (B) The superimposition of the Dengue and Zika NS2B-NS3 proteases in complex with Curcumin with the β-strand colored in purple and loop in green for the NS2B cofactor, as well as the β-strand in brown, helix in cyan and loop in light blue for the NS3 protease domain, but with the Zika protease all colored in blue (I). The structure of the Zika NS2B-NS3 protease in complex with Curcumin in electrostatic potential surface (II).

The binding cavity on the Dengue protease for Curcumin, as well as the binding mode of Curcumin are highly similar to those in the complex of Curcumin with the Zika protease (I of Fig. 3B), which we previously constructed by AutoDock software without any experimental data (21). This finding is very striking, as it implies that the similar cavities for binding allosteric inhibitors likely exist in all flaviviral NS2B-NS3 proteases. Furthermore, the cavity is well recognizable by the current docking programs, thus suitable for the future virtual screening to identify allosteric inhibitors for flaviviral NS2B-NS3 proteases. Nevertheless, the electrostatic surfaces of the cavities of the Dengue and Zika proteases show some difference: while the Dengue one is non-polar and positively-charged (II of Fig. 3A), the Zika one is largely negatively-charged (II of Fig. 3B). This may partly account for the slightly difference of inhibitory effects of Curcumin on flaviviral NS2B-NS3 proteases.

### Molecular dynamics (MD) simulations

Molecular dynamics simulation is a powerful tool to gain insights into protein dynamics underlying protein functions including the enzymatic catalysis (35-39). In particular, it offers insights into the dynamically-driven allosteric mechanisms for the enzyme catalysis. For example, previously by MD simulations in conjunction with the correlation analysis, we successfully revealed that allosteric mechanisms for two mutants of the SARS 3C-like protease are dynamically-driven, because their crystal structures are almost identical to the wild-type. On the other hand, the catalytic activity of one mutant was almost inactivated (38) while that of another was largely enhanced (39).

Therefore, here to understand the mechanism by which Curcumin allosterically inhibits the Dengue NS2B-NS3 protease, we set up three independent MD simulations up to 50 ns respectively for the Dengue NS2B-NS3 protease in the unbound state and in complex with Curcumin. Fig. 4A presents the trajectories of root-mean-square deviations (RMSD) of Cα atoms averaged over three independent simulations of the NS2B and NS3 protease domain of the Dengue protease in the unbound state (blue) and in complex with Curcumin (purple). For NS2B, the averaged RMSD values are 5.43 ± 0.52 and 7.06 ± 0.65 Å respectively for the unbound and bound proteases. For the NS3 protease domain, the averaged RMSD values are 5.58 ± 0.69 and 5.38 ± 0.64 Å respectively for the unbound and bound proteases. This set of the results clearly indicates that the Curcumin binding led to a significant conformational change in NS2B but not in NS3. This observation is in a general agreement with the NMR results that upon binding to Curcumin, the NS2B factor becomes more dynamic than the NS3 protease domain (Fig. 2A and 2B).

**Figure 4.**
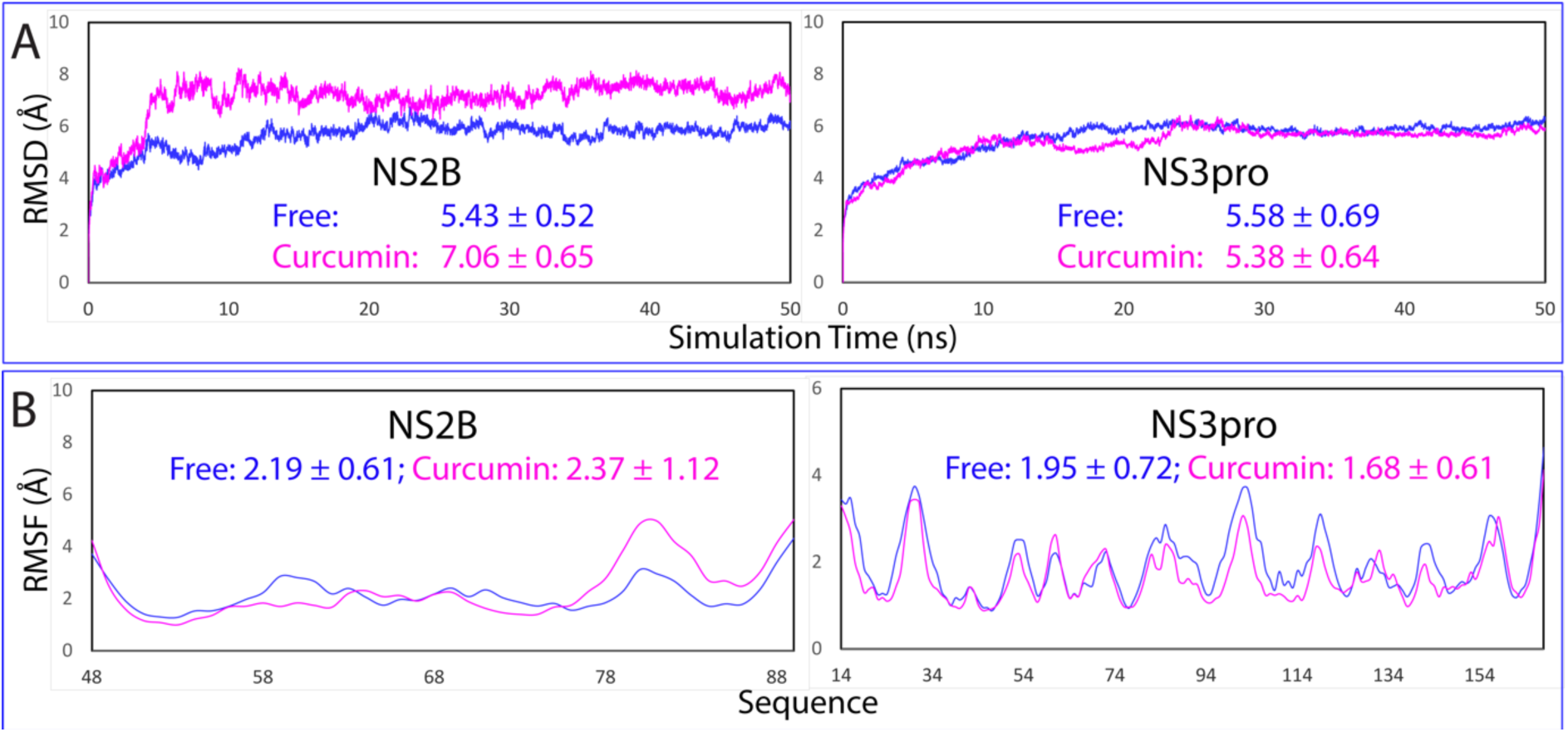
Overall dynamic behaviors revealed by MD simulations. (A) Root-mean-square deviations (RMSD) of the Cα atoms (from their positions in the initial structures used for MD simulations after the energy minimization) averaged over three independent MD simulations of the NS2B and NS3 protease domain respectively. (B) Root-mean-square fluctuations (RMSF) of the Cα atoms averaged over three independent MD simulations of the NS2B and NS3 protease domain respectively.

Similar dynamic behaviours are also reflected by the root-mean-square fluctuations (RMSF) of the Cα atoms in the MD simulations. Fig 4B presents the averaged RMSF of three independent simulations, in which for NS2B, the averaged RMSF values are 2.19 ± 0.61 and 2.37 ± 1.12 Å respectively for the unbound and bound proteases, while for the NS3 protease domain, the averaged RMSD values are 1.95 ± 0.72 and 1.68 ± 0.61 Å respectively for the unbound and bound proteases. The RMSF values indicate that during simulations, the conformational fluctuation of the NS3 protease domain bound with Curcumin becomes even slightly reduced. On the other hand, upon binding to Curcumin, the conformational dynamics of the NS2B cofactor becomes increased. In particular, upon binding to Curcumin, the C-terminal residues Ile76-Glu89 become highly dynamic. However, it is important to point out that the current 50-ns simulations are insufficient to capture the complete transition from the closed (active) to the open (inactive) conformations which occurs over μs-ms time scale.

Fig. 5 presents the structure snapshots in the first sets of MD simulations for the Dengue protease in the unbound state (Fig. 5A) and in complex with Curcumin (Fig. 5B). Consistent with the RMSD and RMSF results (Fig. 4), the NS3 protease domain in the unbound state appears to be slightly more dynamic than that in complex with Curcumin. By contrast, for the NS2B cofactor, the short antiparallel β-hairpin formed over the residues Ser75-Ser79 and Gly82-Ile86, which is characteristic of the closed conformation and has direct contacts with the substrate (II of Fig. 1A), shows dramatic differences in the two simulations. While this β-hairpin retains and only has small fluctuations over its original position in the unbound state in 50-ns simulation (Fig. 5A), its secondary structure becomes unfolded and significantly displaced from its original position in the state complexed with Curcumin (Fig. 5B). More specifically, after 10 ns, the secondary structure of this β-hairpin is already unfolded and moves away from its original position to become highly exposed to the bulk solvent. After 20 ns, a new parallel β-hairpin is formed between the NS2B residues Ile73-Ile76 and NS3 residues Gly114-Lys117, which appears to result in the high exposure of the NS2B residues after Ser79, and may also partly account the observation that the NS3 protease domain is less dynamic in the Curcumin-bound state than in the unbound state. Together, the MD simulation results reveal that the binding of Curcumin disrupts and/or destabilizes the closed conformation which has been well-established to be essential for the catalytic activity, and consequently allosterically inhibits the Dengue NS2B-NS3 protease.

**Figure 5.**
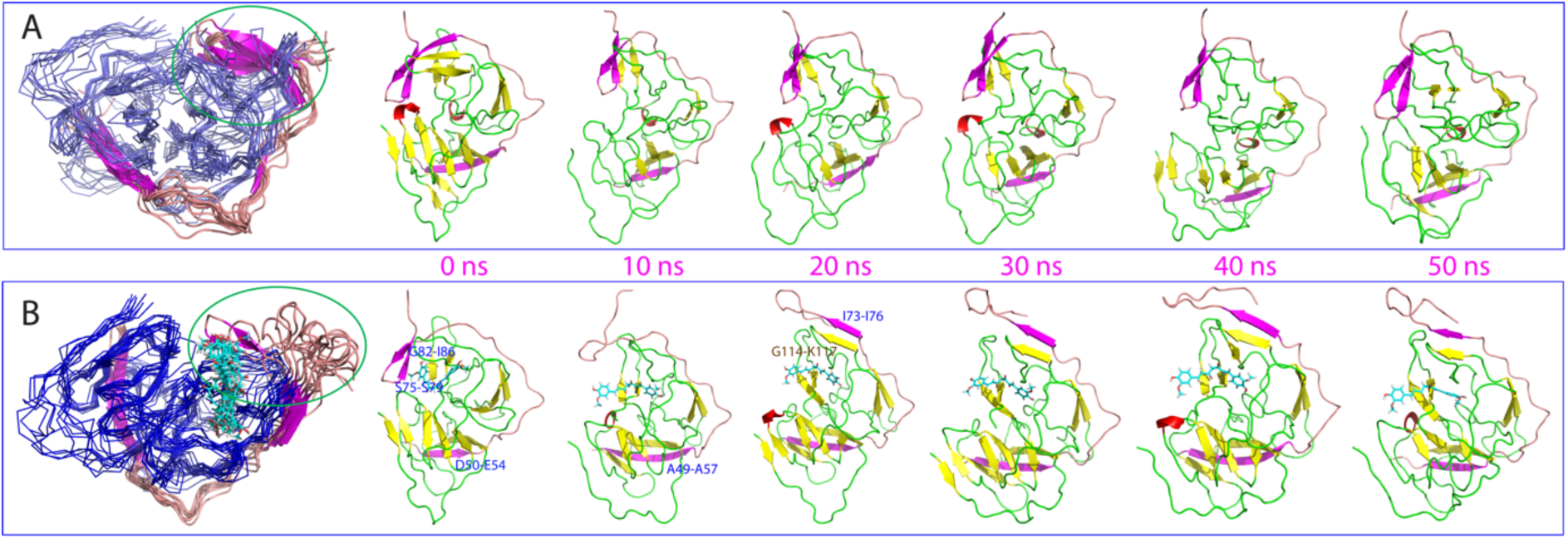
Structure snapshots of MD simulations. (A) Superimposition of 11 structures (one structure for 5-ns interval) of the first set of MD simulation for the Dengue NS2B-NS3 protease in the unbound state as well as 6 individual structures at different simulation time points. (B) Superimposition of 11 structures (one structure for 5-ns interval) of the first set of MD simulation for the Dengue NS2B-NS3 protease in complex with Curcumin, as well as 6 individual structures at different simulation time points. For the clarity of the superimposition, the NS3 protease domain is displayed in the blue cartoon and the NS2B cofactor displayed in ribbon with the β-strand colored in purple and loop in brown. For 6 individual structures, the β-strand is colored in purple and loop in brown for the NS2B cofactor, while the β-strand is in yellow and helix in red and loop in green for the NS3 protease domain. The green circles are used to indicate the NS2B regions forming the short antiparallel β-hairpin characteristic of the closed (active) conformation.

## DISCUSSION AND CONCLUSIONS

The unique NS2B-NS3 serine proteases are highly conserved in Flaviviruses and have been established as a key target for discovery/design of inhibitors to treat Flavivirus infections (4-24,40-43). Currently, most campaigns targeted their active sites for the discovery/design of competitive inhibitors (22,23). However, due to the intrinsic features of their active sites, the design of their active-site inhibitors has been shown to be highly challenging. In this context, one promising solution is to discover/design of their allosteric inhibitors which bind to the cavities other than the active sites. Nevertheless, due to the general challenge in understanding the mechanisms for allosteric processes, only a few attempts were devoted to developing allosteric inhibitors of the Flaviviral proteases (18,19,21,22,40-43). In particular, so far no complete mechanism has been elucidated for the allosteric inhibition on any Flaviviral NS2B-NS3 proteases.

In the present study, by enzymatic kinetic assay, NMR characterization and molecular docking, we have successfully identified a cavity on the Dengue NS2B-NS3 protease which is susceptible to the allosteric inhibition by Curcumin. Together with our previous results with the Zika NS2B-NS3 protease (21), the current study implies that this cavity appears to be conserved in most, if not all, Flaviviral proteases. This cavity is within the allosteric region centred around A125 previously identified (III of Fig. 3A) (18). Furthermore, a synthetic small molecule is also bound to a cavity over this region (Fig. 6A-C) (19). Therefore, these results together suggest that the patches/cavities around this region may represent promising therapeutic targets for further discovery/design of allosteric inhibitors for the Flaviviral NS2B-NS3 proteases.

**Figure 6.**
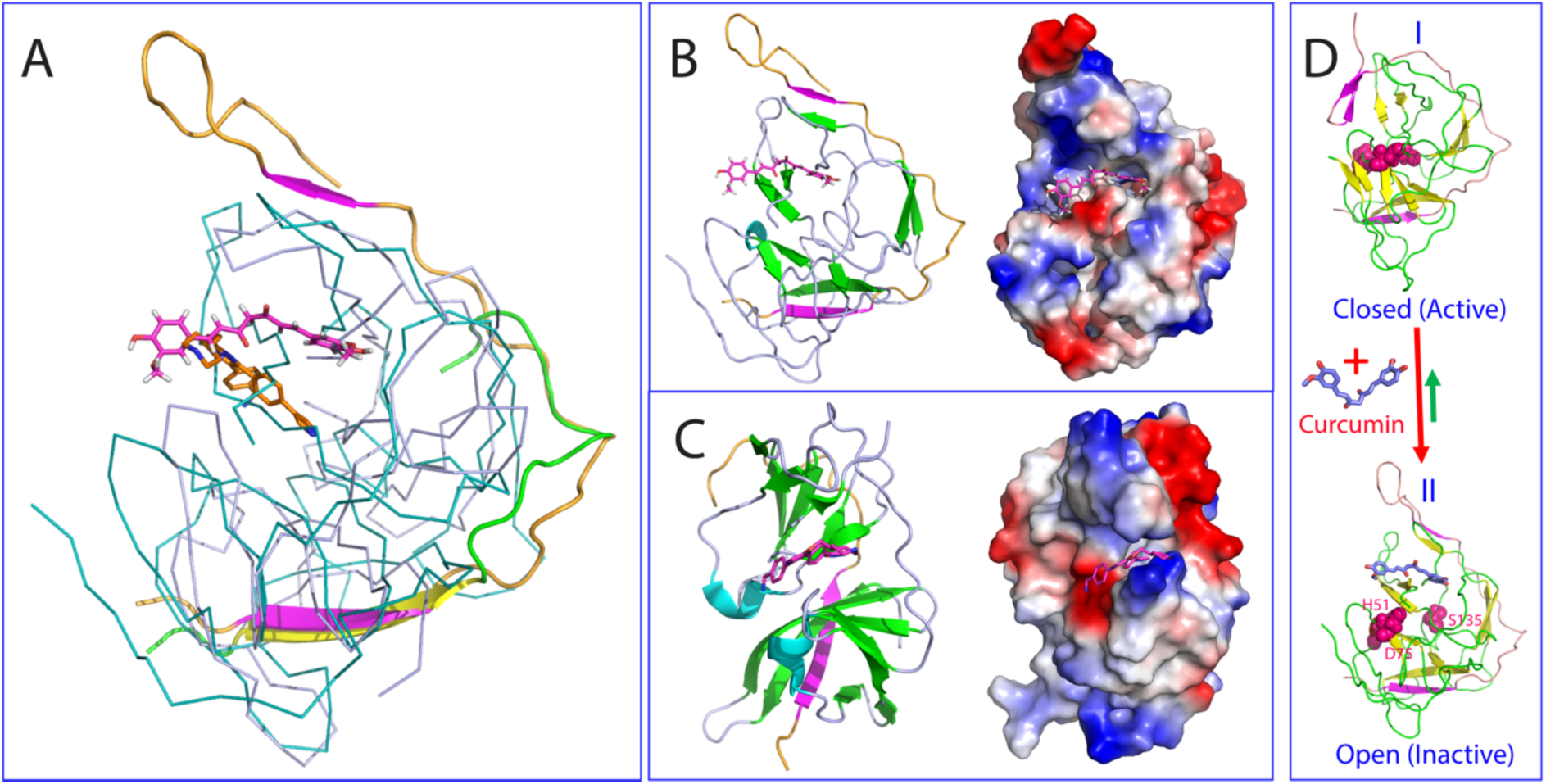
Curcumin allosterically inhibits the Dengue protease by disrupting the active conformation. (A) Superimposition of the simulation structure at 50 ns of the Dengue NS2B-NS3 protease complexed with Curcumin with the NS3 protease domain displayed in the light blue cartoon and the NS2B cofactor displayed in ribbon with the β-strand colored in purple and loop in brown; as well as crystal structure of the Dengue NS2B-NS3 protease complexed with a allosteric inhibitor (PDB ID of 6MO2) with the NS3 protease domain displayed in the cyan cartoon and the NS2B cofactor displayed in ribbon with the β-strand colored in yellow and loop in green. (B) The simulation structure at 50 ns of the Dengue NS2B-NS3 protease complexed with Curcumin in ribbon and in electrostatic potential surface. (C) The crystal structure of the Dengue NS2B-NS3 protease complexed with a synthetic allosteric inhibitor in ribbon and in electrostatic potential surface. (D) The proposed mechanism by which Curcumin allosterically inhibits the Dengue NS2B-NS3 protease through disrupting the closed (active) conformation to modulate the conformational equilibrium.

Due to the general beneficial effects of Curcumin, intensive efforts have been devoted to synthesizing curcuminoids with the improved activity and bioavailability (4,25-27,45). In the further, it is of significant interest to apply our methods established here to characterize the interactions and mechanisms for Curcumin to interact with other target proteins, as well as the interactions of Curcumin analogues including its metabolites with the Flaviviral NS2B-NS3 proteases to obtain better inhibitors.

So a key question arises as how Curcumin achieves the allosteric inhibition on the NS2B-NS3 proteases if it binds to a cavity other than the active site. Previously, our studies on the SARS 3C-like protease revealed two major mechanisms for the allosteric regulation: the structurally-driven and dynamically-driven allostery (24,38,39,44). While the structurally-driven allostery could be characterized by experimentally determining the structures by X-ray crystallography or NMR spectroscopy (44), the dynamically-driven allostery still remains a challenge to be elucidated by classic experimental approaches. Previously, by MD simulations, we successfully decoded the dynamically-driven mechanisms for the opposite allosteric effects of several mutants of the SARS 3C-like proteases, all of which have crystal structures almost identical to that of the wild type (38,39).

Here our NMR characterization reveals that Curcumin induces no detectable change on the averaged structure of the Dengue protease but instead triggers significant dynamics particularly on µs-ms time scale, thus revealing that Curcumin allosterically inhibits the enzyme mainly by dynamically-driven allostery. Indeed, our simulations clearly reveal that the binding of Curcumin has no significant effect on the NS3 protease domain. By contrast, it leads to the unfolding of the short anti-parallel β-hairpin formed by the NS2B residues Ser75-Ser79 and Gly82-Ile86, as well as displacement of the NS2B C-terminus from the original position to become highly exposed (Fig. 5). Therefore, Curcumin appears to achieve the allosteric inhibition by disrupting/destabilizing the close (active) conformation consequently to modulate the conformational equilibrium (Fig. 6D). It has been established that in solution the NS2B-NS3 proteases have two conformations exchanging on µs-ms time scale: in the open (inactive) conformation, NS2B is only partially bound to NS3 and its C-half is far from the active site (I of Fig. 1A), while in the closed (active) state, NS2B is fully tied around NS3 and its C-half folds into an antiparallel β-hairpin which even becomes part of the active site (II of Fig. 1A). Previously we experimentally showed that without the NS2B cofactors, both Dengue and Zika NS3 protease domains have no capacity to correctly fold and thus become highly insoluble (16,21). Furthermore, the truncated NS2B cofactors with the C-half deleted are capable of assisting the folding of the Dengue and Zika NS3 protease domains into the molten-globule like states which are catalytically-inactive and have a large portion of NMR HSQC peaks too broad to be detected, exactly as what we observed here on the Dengue NS2B-NS3 protease in the presence of Curcumin at high concentrations. As such, the binding of Curcumin may induce the population increase of the open (inactive) conformation, as previously observed on the Dengue protease with Ala125 covalently perturbed (18).

Recently, we found that a global network of correlation motions also exists in the SARS 3C-like protease (38,39). An extra-domain mutation N214A far away from the active site is sufficient to decouple the global correlated motions and consequently results in the inactivation of the enzymatic catalysis (38). Here, the correlation analysis of three independent MD simulations for the Dengue NS2B-NS3 protease in the unbound state (Fig. S2A) and in complex with Curcumin (Fig. S2B) clearly revealed that like the SARS 3C-like protease, a global correlation network does also exist in the Dengue NS2B-NS3 protease in the unbound state (Fig. S2A). This global network is largely decoupled upon binding to Curcumin (Fig. S2B), which is similar to what was observed on the inactivated N214A mutant of the SARS 3C-like protease (38). Therefore, although Curcumin binds to a cavity of the Dengue protease with no overlap with the active site and triggers no significant change of the NS3 conformation, the binding is sufficient to globally decouple the network of correlation motions, thus impairing the catalysis of the protease. In the future it is of fundamental interest to continue our MD simulations up to μs or even ms to visualize the transition from the closed into the open conformations induced by the binding of Curcumin.

In conclusion, here by NMR studies and molecular docking, we successfully mapped out the binding cavity of Curcumin onto the Dengue NS2B-NS3 protease, which has no overlap with the active site. Further MD simulations decode that Curcumin achieves allosteric inhibition by disrupting/destabilizing the closed (active) conformation to modulate the conformational equilibrium. With previous results from other and our groups, the current study thus leads to the establishment of a region most likely conserved in all flaviviral NS2B-NS3 proteases, which can serve as a therapeutic target for discovery/design of small molecule allosteric inhibitors. Finally, as it is an extremely daunting task to develop a compound into a marketed drug, Curcumin owns a great advantage in being immediately utilized to fight the flaviviral infections, as well as serves as a promising lead for further design of potent allosteric inhibitors.

## EXPERIMENTAL SECTION

### Plasmid construction, protein expression and purification

In this study, we used the expression plasmids constructed in our previous studies (16), which encodes DENV2 NS3 (14-185) and NS2B (48-100) with the same starting and ending residues as the constructs used in the previously published NMR study (14). The recombinant proteins of NS2B and NS3 were expressed in *Escherichia coli* BL21 (DE3) Star cells following the exactly the same protocol we established before (16). The generation of the isotope-labeled proteins followed the same procedure except for the growth of bacteria in M9 medium with the addition of (^15^NH_4_)_2_SO_4_ for ^15^N labeling and (^15^NH_4_)_2_SO_4_/[^13^C]-glucose for ^15^N-/^13^C-double labelling (16). The purity was checked by SDS-PAGE gels and molecular weights were verified by ESI-MS and Voyager STR time-of-flight-mass spectrometer (Applied Biosystems). The protein concentration was determined by the UV spectroscopic method in the presence of 8 M urea (46).

### Enzymatic kinetic assay

Enzymatic kinetic assay of the Dengue NS2B-NS3 protease followed the same protocol we previously described for the Dengue and Zika NS2B-NS3 proteases (16,21). All enzymatic assays were performed in triplicates. Substrate peptide was Bz-Nle-Lys-Arg-Arg-AMC (Bz-nKRR-AMC), which was purchased from GenScript (Piscataway, NJ), while HPLC-purified Curcumin was from Sigma Aldrich with the purity >93%. Stock solution of curcumin was dissolved in DMSO. The protease is in 50 mM Tris-HCl (pH 7.5), 0.001% Triton X-100, 0.5 mM EGTA at 37°C. To determine IC50 for Curcumin, 50 nM protease was incubated with various concentrations of Curcumin (in 1μl DMSO) at 37 °C for 30 mins, and Bz-nKRR-AMC addition to 250 µM initiated enzymatic reaction. To determine Ki for Curcumin, the same kinetic assay was performed with different final concentration of Bz-nKRR-AMC and Curcumin. Enzymatic reaction was monitored with fluorescence upon hydrolysis of substrate peptide (Bz-nKRR-AMC) at λex of 380 nm and λem of 450 nm. Fluorescence values (relative fluorescence units/sec) were fitted to the non-competitive inhibition model in GraphPad Prism 7.0; Ki was obtained with fitting to equation: Vmaxinh = Vmax/(1+I/Ki), while I is the concentration of inhibitor (16,21).

### NMR characterization of the binding

2D ^1^H-^15^N HSQC NMR experiments were acquired on an 800 MHz Bruker Avance spectrometer equipped with pulse field gradient units as described previously (16,21). NMR samples of 100 μM protease were prepared in 10 mM phosphate buffer, pH 7.5, 5% DMSO and 10% D_2_O for NMR spin-lock. All NMR experiments were done at 25 °C. Curcumin was also dissolved in the same buffer.

### Molecular docking

Here the crystal structures of the Dengue NS2B-NS3 protease in the closed form (PDB code: 3U1I) (10) and in the open conformation (PDB ID of 4M9T) (18) were used for docking. Chemical structure of Curcumin was downloaded from ChemicalBook database (http://www.chemicalbook.com), and their structural geometry were generated and optimized with Avogadro (47). NMR titration derived constraints were used to guide the docking by HADDOCK software (32) and CNS (48). CNS topology and force field parameters of Curcumin is converted from PRODRG server (49). The docking of the Curcumin-protease complex was performed in three stages: (1) randomization and rigid body docking; (2) semi-flexible simulated annealing; and (3) flexible explicit solvent refinement, as we extensively performed (33,34). The complex structures with the lowest energy score were selected for the detailed analysis and display by Pymol (PyMOL Molecular Graphics System, Version 0.99rc6 Schrödinger, LLC).

### Molecular dynamics (MD) simulations

The crystal structure of the Dengue NS2B-NS3 protease (PDB code: 3U1I) in the closed conformation was used as the unbound form while the docking structure of the Curcumin-protease complex with the lowest energy was used as the bound form in the current MD simulations with three independent simulations for each of them. Electrostatic potential of both Curcumin was first calculated with the 6-31G(d,p) basis set using the GAUSSIAN 16 program, which is converted into partial charge of individual atoms using restrained electrostatic potential (RESP) procedure in Antechamber program (50). The topology parameters of Curcumin were obtained using GAFF (51). All MD simulations reaching 50 ns were conducted using GROMACS (52), and AMBER99SB-IDLN all-atom force field (53) parameters.

The simulation system is a periodic cubic box with a minimum distance of 12 Å between the protein and the box walls to ensure the proteases does not interact with its own periodic images during MD simulation. About 13,000 water molecules (TIP3P model) solvated the cubic box for the all atom MD simulation. 2 Na^+^ ions were randomly placed to neutralize the charge of protein complex. The long-range electrostatic interactions were treated using the fast particle-mesh Ewald summation method (54). The time step was set as 2 fs. All bond lengths including hydrogen atoms were constrained by the LINCS algorithm (55). Prior to MD simulations, the initial structures were relaxed by 500 steps of energy minimization, followed by 100 ps equilibration with a harmonic restraint potential.

### Correlation analysis

To analyze correlated motions for residues of the complex, we utilize MutInf approach, which calculates correlated motion between pairs of residues using low-frequency motions (internal coordinates) that are represented by dihedral angles (56). The approach here uses internal coordinates of different protein conformers obtained from MD simulations, and converts them into configurational entropy expansion terms which are computed in an entropy-based correlation matrix. The degree of correlated motions between different pairs of residues with correlated conformations is represented by a metric called mutual information (MutInf) (56). In this present study, we first normalized MutInf values and 0.55 was were used as the threshold value to define pairs of residues as highly correlated motions.

## Acknowledgement

This study is supported by Ministry of Education of Singapore (MOE) Tier 1 Grant R-154-000-B45-114 to Jianxing Song.

